# Accurate Reference-Free Somatic Variant-Calling by Integrating Genomic, Sequencing and Population Data

**DOI:** 10.1101/383703

**Authors:** Ren X. Sun, Christopher M. Lalansingh, Shadrielle Melijah G. Espiritu, Cindy Q. Yao, Takafumi N. Yamaguchi, Stephenie D. Prokopec, Lesia Szyca, Kathleen E. Houlahan, Lawrence E. Heisler, Morgan Black, Constance H. Li, John W. Barrett, Anthony C. Nichols, Paul C. Boutros

## Abstract

The detection of somatic single nucleotide variants (SNVs) is critical in both research and clinical applications. Studies of human cancer typically use matched normal (reference) samples from a distant tissue to increase SNV prediction accuracy. This process both doubles sequencing costs and poses challenges when reference samples are not readily available, such as for many cell-lines. To address these challenges, we created S22S: an approach for the prediction of somatic mutations without need for matched reference tissue. S22S takes underlying sequence data, augments them with genomic background context and population frequency information, and classifies SNVs as somatic or non-somatic. We validated S22S using primary tumor/normal pairs from four tumor types, spanning two different sequencing technologies. S22S robustly identifies somatic SNVs, with the area under the precision recall curve reaching 0.97 in kidney clear cell carcinoma, comparable to the best tumor/normal analysis pipelines. S22S is freely available at http://labs.oicr.on.ca/Boutros-lab/software/s22s.

---

Technological advances in DNA sequencing have enabled routine analysis of cancer genomes. The identification of somatic single nucleotide variants (SNVs) is commonly used to catalog mutational landscapes^1–3^, to link genomics with drug efficacy^4–6^ and to create clinically useful diagnostics^7,8^. In most of these studies, predictions are made from matched tumor/normal pairs, and large benchmarks have evaluated their accuracy^9–12^. There is, however, an increasing number of applications where matched normal reference samples are not readily available. Normal samples comprise about half of sequencing costs in clinical studies, and their collection is not always routine or even possible for retrospective cohorts that include deceased patients. Similarly, cell-line studies, such as those used for drug-screening, rarely have matched normal samples.

However SNV identification without a reference sample significantly reduces detection accuracy^13–15^. Many groups resort to *ad hoc* metrics or repurposing tools originally designed for germline analysis^4^’;;^15,16^, since only a small number of analytical tools accommodate unmatched tumor samples. The most popular approach is to generate a surrogate normal from a pool of normal samples^17–19^, which only removes false positives resulting from germline contamination, not other types of errors^15–17^. The resulting datasets anecdotally have high false-positive rates, but no systematic benchmark yet exists^13^. To address this challenge, we created **S**ingle-sample **S**omatic **S**NV **S**elector (S22S): a random forest classifier that acts to identify true somatic SNVs from single-sample tumor sequencing data. S22S classifies SNVs as somatic or not by integrating sequencing characteristics, background genomic context (such as GC content, homopolymer rate and trinucleotide context) and population prevalence of both germline and somatic mutations (see **Online Methods**).

To generate a set of gold-standard for model training, we used samples with matched-normal references from three tumor types from the Cancer Genome Atlas (TCGA) network-head and neck squamous cell carcinoma (HNSC)^20^, kidney renal clear-cell carcinoma (KIRC)^21^, prostate adenocarcinoma (PRAD)^22^-and a breast cancer dataset that utilized a targeted sequencing platform^23^.S22S operates under the hypothesis that true somatic calls possess distinct profiles of sequencing properties, genomic contexts and population frequencies and thus can be discriminated from false calls. To build the feature set, SNVs were called for each sample in four ways: (1) the normal sample was used to identify germline variants using GATK^24–26^, (2) somatic mutations were identified by an SNV caller (MuTect^17^, SomaticSniper^27^ or Strelka^28^) using the tumour-normal pair and (3) GATK (HaplotypeCaller) and (4) MuTect^17^ with a panel of normal samples (PoN) were used to call SNVs from tumor samples alone to capture characteristics of unmatched analyses (see **Online Methods**). To facilitate appropriate model testing, 30% of the samples from each tumor type were set aside for validation. A separate PoN was generated for each tumor type – BRCA, HNSC, KIRC and PRAD – by running MuTect on each normal sample from the training set individually and subsequently aggregating the results. SNVs were then assigned true or false class labels according to the overlap between the four call-sets, where all somatic SNVs detected from T/N analysis were deemed as true positives and all other genomic positions as true negatives (**Figure 1A**). To optimize signal for machine-learning, in the training set those (rare) calls that were both somatic (calls predicted to be somatic by an SNV caller on T/N pair) and germline (GATK on reference samples) were removed, but these were retained in the validation set to avoid bias in assessment. We used somatic SNV calls from three different algorithms (MuTect^17^, SomaticSniper^27^ or Strelka^28^), as well as the intersecting ensemble set of calls, to facilitate labeling of the true class to avoid model bias and overfitting to any particular tool. The overall process is outlined in **Figure 1A**.

**Figure 1.**
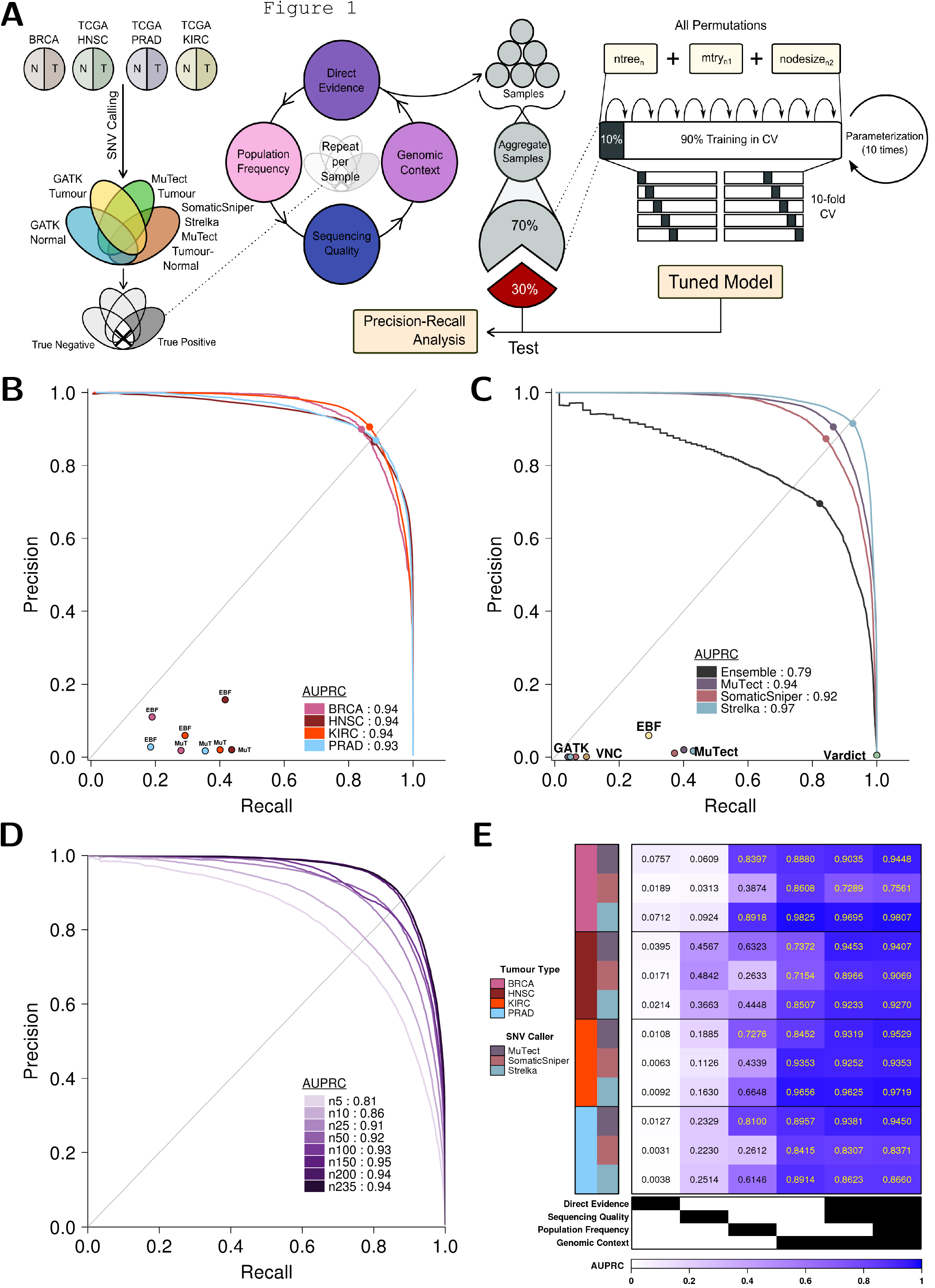
S22S predicts somatic variants without reference samples. A schematic of the algorithm is shown in (A). Per sample, S22S extracts features from the VCF file and from the tumor BAM file and annotates these with population frequency data from publicly available germline and somatic databases to build a feature vector for that sample. We set aside 30% of the samples for validation as an independent test set and aggregate the feature matrices for the remaining 70% of the samples to generate our training set. We used 10-fold cross validation on the training set to parameterize our model. The optimal parameters are used to train a random forest on the full training set. Finally, we assessed model performance on the held-out test set. This procedure was repeated for all four tumor types. Overall model performance for models trained using somatic calls from MuTect is shown in **(B)**. Each model performed well in cis-tumor application on the test dataset, with an AUPRC exceeding 0.9 in all cases. This is superior to existing reference-free SNV calling methods such as EBFilter (EBF) and MuTect using a PoN (MuT). Model performance remains high with other SNV callers **(C)**. The models trained on somatic calls from several different SNV callers for the KIRC dataset are shown, where the AUPRC reaches 0.97 when Strelka is used, and far exceeds the performance of other reference-free somatic classification methods, EBFilter (EBF), VarDict, Virtual Normal Correction (VNC) and MuTect using a PoN (MuTect). S22S is robust to decreased sample-size **(D)**. PR-curves are constructed from models trained on subsets of the full training data for KIRC using somatic calls from MuTect. The lower AUPRC seen with smaller sample sizes demonstrates that adding additional samples will increase the predictive power of the final model. Further investigation of how model decisions are made reveals that certain subsets of features are more predictive of somatic status, but that a model trained using the full set of features had highest performance overall **(E)**.

To accurately identify somatic SNVs in the absence of a normal sample, we leveraged four largely orthogonal data types. First, we incorporated the frequency of known mutations – both at the gene and variant levels – in both normal populations and tumor cohorts (where available). Second, we included metrics indicating sequence quality, such as mapping quality, coverage and position along the read. Third, the genomic context of each variant was considered, including elements like homopolymer rate and GC content, which are correlated with errors in somatic SNV prediction^9^, and trinucleotide context, to adjust for trinucleotide mutational signatures^29^. Fourth, direct evidence supporting the mutation was measured, for example with the number of reads supporting each allele. **Table 1** lists all features. These were used for hyper-parameter tuning *via* 10-fold cross-validation and model fitting in the training cohort (Figure 1A).

**Table 1.** Classification Features. We extracted a total of 59 features for PRAD and 61 features for BRCA, KIRC and HNSC. Features are categorized based on origin and purpose. The PRAD model has two fewer features than the other three models (ClippingRankSum and SOR are missing), as it was analyzed using an older version of GATK, which lacked these two metrics.

| FEATURE ALIAS | FEATURE DESCRIPTION |
| --- | --- |
| **Population Frequency** |  |
| AF.1000Genomes | Global allele frequency based on AC/AN |
| AFR_AF | Allele frequency for the Afriacn population based on AC/AN from 1000 Genomes Project |
| AMR_AF | Allele frequency for the American population based on AC/AN from 1000 Genomes Project |
| ASN_AF | Allele frequency for the Asian population based on AC/AN from 1000 Genomes Project |
| EUR_AF | Allele frequency for the European population based on AC/AN from 1000 Genomes Project |
| LDAF | Allele frequency as inferred from haplotype estimation |
| NHLBI_EA_MAF | Minor allele frequency in percent for European population from NHLBI |
| NHLBI_AA_MAF | Minor allele frequency in percent for African American population from NHLBI |
| NHLBI_ALL_MAF | Overall minor allele frequency from NHLBI |
| TCGA.fraction.MAF | Proportion of recurrence in the TCGA dataset, position-based |
| TCGA.fraction.RecSNV | Proportion of recurrence in the TCGA dataset, gene-based |
| **Sequencing Quality** |  |
| BaseQRankSum | Z-score from Wilcoxon rank sum test of ALT Vs. REF base qualities |
| BaseQual | Mapping quality scores (in phred-scale) for probability of misidentified base |
| ClippingRankSum | Z-score From Wilcoxon rank sum test of Alt vs. Ref number of hard clipped bases |
| DistanceIndel | Distance (in bases) to closest known germline Indel |
| DistanceSNP | Distance (in bases) to closest known germline SNP |
| FS | Phred-scaled p-value using Fisher's exact test to detect strand bias |
| MappingQual | Mapping quality scores (in phred-scale) for probability of misplaced read |
| MLEAC | Maximum likelihood expectation (MLE) for the allele counts |
| MLEAF | Maximum likelihood expectation (MLE) for the allele frequency |
| MQ | Root Mean-squared (RMS) Mapping Quality |
| MQRankSum | Z-score From Wilcoxon rank sum test of ALT vs. REF read mapping qualities |
| NonRefForward | Quality of at least 13 non-REF bases on forward strand |
| NonRefReverse | Quality of at least 13 non-REF bases on reverse strand |
| QD | Variant Confidence/Quality by Depth |
| ReadPosRankSum | Z-score from Wilcoxon rank sum test of ALT vs. REF read position bias |
| ReadPosition | Position within the read |
| RefForward | Quality of at least 13 REF bases on forward strand |
| RefReverse | Quality of at least 13 REF bases on reverse strand |
| StrandBias | Proportion of reads mapped to forward strand |
| SOR | Symmetric Odds Ratio of 2x2 contingency table to detect strand bias |
| SumNonRefBase | Sum of non-ref base qualities |
| SumNonRefMap | Sum of non-ref mapping qualities |
| SumRefBase | Sum of reference base qualities |
| SumRefMap | Sum of ref mapping qualities |
| SumSqNonRefBase | Sum of squares of non-ref base qualities |
| SumSqNonRefMap | Sum of squares of non-ref mapping qualities |
| SumSqRefBase | Sum of squares of reference base qualities |
| SumSqRefMap | Sum of squares of ref mapping qualities |
| SumSqTailNonRef | Sum of squares of tail distance for non-ref |
| SumSqTailRef | Sum of squares of tail distance for ref bases |
| SumTailNonRef | Sum of tail distance for non-ref bases |
| SumTailRef | Sum of tail distance for ref bases |
| TumourCoverage | Read depth from tumour bam file |
| VDB | Variant distance bias for indication of potential misalignment due to nearby SNP |
| **Genomic Context** |  |
| homopolymer.rate | Frequency of homopolymers within 100 bases up and downstream of position |
| GC | Proportion of bases that are GC within 100 bases up and downstream of position |
| trinucleotide | Trinucleotide context of position |
| **Direct Evidence** |  |
| AC | Allele count for ALT allele |
| AF | Allele frequency for ALT allele |
| ALT.AD | Allelic depth for ALT |
| ALT.AF | Calculated allelic frequency of ALT (ALT.AD/DP) |
| AN | Total number of alleles called in genotype |
| DP | Approximate read depth, VCF-derived |
| DP.pileup | Number of reads covering or bridging position |
| gatk | Presence of call using GATK in tumour-only mode |
| mutect | Presence of call using MuTect with a panel of normals |
| NonRefAllele | Forward non-ref allele |
| REF.AD | Allelic depth for REF |
| REF.AF | Calculated allelic frequency of REF (REF.AD/DP) |
| RefAllele | Forward ref allele |

To evaluate the performance of S22S, we first used the respective held-out test datasets for each tumor (n_BRCA_ = 164, n_HNSC_ = 113, n_KiRC_ = 100, n_PRAD_ = 56) and measured the Area <u>U</u>nder the <u>P</u>recision <u>R</u>ecall <u>C</u>urve (AUPRC). In all four test datasets, S22S showed high precision and recall (**Figure 1B**). We selected our operating point for each dataset (*i.e.* the threshold used for binary classification, representing a specific precision-recall tradeoff) to be that with the maximum F_1_ score. To compare the performance of S22S with alternative approaches, we examined the precision and recall of five different tumor-only approaches. We first considered the use of a germline caller (GATK) on the tumor sample. Second, we looked at the use of MuTect on the tumor sample using a surrogate reference normal cohort derived from the training dataset (in PoN mode). We also considered three additional reference-free SNV prediction tools: Virtual Normal Correction (VNC)^18^, EBFilter^19^ and VarDict^30^. None of these exceeded a precision of 0.10 for any operating point in either of the four tumors assessed, and S22S was Pareto superior to all of them (**Figure 1B-C, Supplementary Figure 1, Supplementary Data 1**).

To demonstrate the impact of improved reference-free somatic classification on applications evaluating mutational landscapes, we looked at the ten most recurrently-mutated genes reported by the original studies^20–23^. For each recurrent gene, we calculated the proportion of correctly predicted mutations (true positive rate) and the proportion of correctly predicted non-mutated genes (true negative rate). S22S dramatically improved true negative rates for all recurrent genes (**Supplementary Figure 2**). To extend its application, we assessed the ability of S22S to identify recurrently-mutated genes in cell line studies and applied the HNSC model trained on MuTect calls to a handful HNSC cell lines and compared the results with a commonly-used surrogate – MuTect with a PoN (**Supplementary Figure 3**). We compared the top 20 recurrent genes identified by both methods – S22S with minimal filtering (dbSNP and COSMIC only) and MuTect with a PoN with extensive filtering (see **Somatic SNV Calling** in **Online Methods**). While the results from using MuTect with a PoN was nearly uninterpretable – many genes identified as being mutated across all cell lines, thus only the top 20 sorted alphabetically are shown – S22S was able to drastically reduce noise and produce some potentially biologically-relevant results.

While the use of TCGA and other large publicly-available datasets provided sufficiently-sized cohorts for method benchmarking, smaller studies (*e.g.* rare tumor types) will have fewer training data available. To assess the sensitivity of S22S performance to training set size, we performed a subset analysis. S22S hyper-parameter optimization and modeling was repeated in a titration series with subsets of the training dataset and validated on the same independent held-out dataset used previously. We used sample sets of 5, 10, 25, 50 and 100 (and increasing additional sets in increments of 50 as sample sizes allowed) for each tumor and trained three models per tumor (one for each SNV caller). For all models, AUPRC increased in large steps up to 50 sample training cohorts, with smaller gains from additional samples beyond this threshold (**Figure 1D, Supplementary Figures 4-7**).

S22S has been shown to improve prediction accuracy in the presence of a training dataset related to the given experiment. However, there are many situations where disease-specific or tool-specific training data is unavailable. In this scenario, a surrogate dataset may prove sufficient. To assess the performance of S22S in such cases, we first applied the model trained on somatic calls from a given SNV caller (*i.e*. MuTect) from one tumor to the test sets of the other three tumors that also used somatic calls from this caller (*i.e*. MuTect), and assessed the cross-tumor model performance (**Supplementary Figure 8**). We next, for each tumor type, applied models trained using different SNV callers to the test sets of the other SNV callers to evaluate the cross-caller model performance (**Supplementary Figure 9**). As expected, performance was decreased by training on a different tumor type or SNV caller, but the AUPRC remained higher than any other method, reaching an AUPRC of 0.90 for the KIRC-trained model (using MuTect) evaluated on the HNSC testing dataset (**Supplementary Figures 8**). This suggests that a similar set of features is important for prediction performance across tumor types as well as SNV calling algorithms.

To the sensitivity of S22S to model features, we conducted a titration study of model features, similar to our sample-size analysis. We repeated model training using a subset of the features each time – with features grouped based on their respective data types (*i.e*. sequencing quality, population frequency, *etc.*, see **Table 1**) – and tested the performance on the respective held-out test sets. We observed that certain categories of features were more predictive of somatic status than others, and that the ranking of categories differed slightly across tumors and SNV calling algorithms, but overall, the best performing models were those that incorporated all features (**Figure 1E**). To assess the contributions of each feature more deeply, we next evaluated the most discriminative features in each model by quantifying the mean decrease in the Gini coefficient. This is a metric that describes node purity and quantifies a variable’s usefulness in partitioning observations^31^. There was a set of features was important both across tumor types as well as SNV callers, as well as some more specific to specific models – this variability in ranking likely accounts for differential performance across tumor types and SNV callers (**Supplementary Figure 10-11**).

In summary, we have demonstrated accurate prediction of somatic variants without matched reference samples by integrating sequencing data with genomic and population prior knowledge. By using these effectively as a prior, S22S outperforms current standards of tumor-only SNV calling in all performance metrics, even when little or no training data are available. Additional development and community efforts can help build a collection of models that are robust and applicable to different tumors, even those with significant contamination such as acute myeloid leukemia. S22S is agnostic to sequencing platform, and can work with WGS, exome or targeted sequencing panels, and its implementation is freely available. Many somatic SNV callers (both tumor/normal and reference-free) incorporate panels of reference samples, but the distribution of such panels can be restricted by their incorporation of germline SNP data. In addition, these normal panels can grow burdensome and costly, both in terms of storage and computations. By contrast, S22S models can be readily shared, and indeed those generated here are available along with the respective training and testing data (**Supplementary Data 2-9**). Large consortia like ExAC, TCGA, ICGC and ICGC-ARGO are ideally placed to generate, in a centralized way, robust models for somatic SNV detection that can be shared with the community as a whole.

## ACKNOWLEDGEMENTS

This work was supported by the Ontario Institute for Cancer Research to P.C.B. through funding provided by the Government of Ontario as well as by Prostate Cancer Canada and is proudly funded by the Movember Foundation, grant #RS2014-01. P.C.B. is supported by a CIHR New Investigator Award and by a Terry Fox Research Institute New Investigator Award. This project was supported by Genome Canada through a Large-Scale Applied Project contract to P.C.B., Sohrab Shah and Ryan Morin. This work was supported by the Discovery Frontiers: Advancing Big Data Science in Genomics Research program, which is jointly funded by the Natural Sciences and Engineering Research Council (NSERC) of Canada, the Canadian Institutes of Health Research (CIHR), Genome Canada, and the Canada Foundation for Innovation (CFI). R.X.S. was supported by Ontario Graduate Scholarship through funding provided by the Government of Ontario and a Graduate Fellowship award through funding provided by the University of Toronto. P.C.B. was supported by a CIHR Project Grant and the Canadian Cancer Society (grant #705649). The study sponsors had no role in the study design; in the collection, analysis and interpretation of data; in the writing of the report; or in the decision to submit the paper for publication. The authors thank X. Lin and V. Sabelnykova for their insights on machine learning and statistics, J.M. MacDonald for development of the HPCI Perl package and A.P. Masella for aid in the filtering framework for SNVs. The authors thank all members of the Boutros Lab for helpful suggestions. The results described here are based in part upon data generated by the TCGA Research Network: http://cancergenome.nih.gov/.

## AUTHOR CONTRIBUTIONS

R.X.S., M.B., J.B., A.C.N. and P.C.B. initiated the project. R.X.S., L.E.H., S.M.G.E. and C.Q.Y. processed TCGA HNSC samples. R.X.S., L.E.H and T.N.Y. processed TCGA PRAD samples. R.X.S and S.D.P processed TCGA KIRC samples. R.X.S. and C.H.L. performed analyses on the sequencing data. R.X.S. was responsible for modeling of the sequencing data. R.X.S., C.M.L, K.E.H. and S.D.P were responsible for pipeline and software development. R.X.S and L.S. were responsible for data visualization. Research was supervised by A.C.N. and P.C.B., and R.X.S. wrote the manuscript, which was edited and approved by all authors.

## ONLINE METHODS

### Datasets

To build the training and test sets for our models, we leveraged publicly-available exome sequencing samples of primary tumors and their matched normal samples from three cancer types published by the The Cancer Genome Atlas (TCGA) network: head and neck cancer (HNSC)^20^, kidney renal clear cell cancer (KIRC)^21^ and prostate cancer (PRAD)^22^, and one tumor type – breast cancer (BRCA) – from a targeted sequencing panel^23^. For TCGA datasets, BAM files were obtained from the Cancer Genomics Hub (CGHub, https://cghub.ucscedu/)^32^ and realigned using BWA^33^ and recalibrated using GATK^24–26^ prior to downstream analyses (described in depth below). Our final HNSC cohort is composed of 376 tumor/normal pairs, our KIRC cohort of 335 tumor/normal pairs and our PRAD cohort is composed of 188 tumor/normal pairs. For the targeted breast cancer dataset, BAM files were obtained from the European Genome-phenome Archive (EGA)^34^, under the ID of EGAD00001002115, and recalibrated using GATK prior to downstream analysis. The final BRCA cohort is composed of 546 tumor/normal pairs. All models trained are available in **Supplementary Data 2-5** while the training and testing matrices for all model building used for the four tumor types are available in **Supplementary Data 6-9**.

### Defining True/False Classes

In order to train our model to accurately recognize true calls from false ones, we took a comprehensive approach when defining our truth and false sets. To create an adequate feature space, we performed SNV calling (focusing on substitutions rather than indels) in four manners. First, we called germline and somatic SNVs using the standard approach with tumor/normal pairs, and used GATK^24–26^ for germline SNPs and and three SNV callers – MuTect, SomaticSniper^27^ and Strelka – for somatic SNVs. We also performed tumor-only SNV calling in two ways, first using GATK in tumor-only mode and subsequently, using MuTect^17^ with a panel of normal samples (PoN) generated from the normal samples of each respective tumor type. We defined the true class using variants identified from an SNV caller on paired samples that were absent in the GATK germline calls. The false class encompassed all calls that were present in the germline set as well as both tumor-only call sets, but not found in the somatic set (**Figure 1A**). To eliminate ambiguity for our learner, we elected to omit any overlaps found between the germline and somatic calls from downstream analysis. We also trained models using somatic calls from three SNV callers to label our truth set in order to avoid learning patterns of any particular SNV calling algorithm.

### Realignment and Recalibration of Sequencing Data

For each TCGA BAM file obtained from CGHub, back-conversion to FASTQ files was done to allow realignment to the human reference genome for standardization, using the SamToFastq function from picard (v.1.92) (http://broadinstitute.github.io/picard). For PRAD, lane-level raw sequencing reads were realigned to human reference build hg19 using bwa^33^ aln (v0.5.7), while for HNSC and KIRC, the realignment was performed with the human reference build GRCh37 with decoy (hs37d5) and bwa mem (v0.7.12). Merging of lane-level BAMs from the same library within each sample was facilitated via picard (v1.92), with duplicates marked, and was followed by library-level merging of BAMs, without marking of duplicates. Quality control metrics were used to assess the coverage for BAM files obtained from EGA as part of the BRCA dataset, and the reported targeted regions all had sufficient coverage, as previously reported^23^. BAM files as part of the BRCA dataset were not realigned prior to recalibration.

We used GATK (v2.4.9 for PRAD samples, v3.4.0 for HNSC samples and v.3.5.0 for BRCA and KIRC samples, the version discrepancy a result of time lapse between sample processing) to perform local realignment and base quality recalibration on each realigned TCGA tumor/normal BAM pair and BRCA BAM file obtained from EGA. Separate tumor and normal sample-level BAM files were extracted, followed by header correction using samtools (v0.1.19) and indexing using picard (v1.107) to generate the final realigned and recalibrated BAM file per sample.

### Germline SNP Calling

We used GATK (v2.4.9 for PRAD, v3.4.0 for HNSC and v3.5.0 for BRCA and KIRC) to call germline single nucleotide polymorphisms (SNPs). For each PRAD sample, we used UnifiedGenotyper, followed by VariantRecalibrator and ApplyRecalibration on the realigned and recalibrated tumor/normal pair and removed all insertions/deletions (INDELs), somatic SNVs and ambiguous SNVs with more than one alternate base to obtain our final germline VCF callset. For BRCA, KIRC and HNSC samples, we used HaplotypeCaller followed by VariantFiltration to hard-filter our callset using the following filter expressions: “QD < 10.0 || FS > 60.0 || MQ < 40.0 || DP < 50 || SOR > 4.0 || ReadPosRankSum < −8.0 || MQRankSum < −12.5” and “MQ0 >= 4 && ((MQ0 / (1.0 * DP)) > 0.1)” to generate the final germline calls. We referred to the GATK Best Practices recommendations for the development of this pipeline^24,26^.

### Somatic SNV Calling

First, we predicted somatic SNVs for all tumor types using three callers, MuTect (v1.1.4 for PRAD and v1.1.6 for HNSC and v1.1.7 for BRCA and KIRC), SomaticSniper^27^ (v1.0.2 for PRAD, v1.0.4 for HNSC and v1.0.5 for BRCA and KIRC) and Strelka (v1.0.12 for PRAD and HNSC and 1.0.14 for BRCA and KIRC) with the tumor/normal pair. All tools were run with default options unless otherwise specified. MuTect was given the same target capture region file as GATK and executed with dbSNP (v138) and COSMIC (v66). SomaticSniper was run with the −q option (mapping quality threshold) set to 1 and then filtered using a series of Perl scripts provided by the SomaticSniper package to remove possible false positives (http://gmt.genome.wustl.edu/packages/somatic-sniper/documentation.html).

Following SNV calling, we applied an additional filtering step where we removed SNVs that were found in any of the following databases (also referred to as “blacklists”) using tabix to produce the final set of somatic calls^35^: dbSNP142^36^ (modified to remove somatic and clinical variants, with variants with the following flags excluded: SAO = 2/3, PM, CDA, TPA, MUT and OM), 1000 Genomes Project (v3)^37^, Complete Genomics 69 whole genomes^38^, duplicate gene database (v68)^39^, ENCODE DAC and Duke Mapability Consensus Excludable databases^40^ (comprising poorly mapping reads, repeat regions and mitochondrial and ribosomal DNA), invalidated somatic SNVs from 68 human colorectal cancer exomes (unpublished data) using the AccuSNP platform (Roche NimbleGen), germline SNPs from 477 samples in previous work in prostate cancer with an additional 10 WGS from prostate cancer patients with high Gleason score^41^ and the Fuentes database of likely false positive variants^42^; SNVs were “whitelisted” (retained independent of presence in other databases) if found in the Catalogue of Somatic Mutations in Cancer (COSMIC)^43^ database (v71).

### Tumor-only SNV Calling

To capture the error profiles of SNVs called in situations where a normal sample was not used, we also performed somatic SNV calling using only the tumor samples. We used two analytical approaches that were most frequently employed in this situation, as seen and discussed across various sequencing forums. The first approach was to run variant calling with GATK in tumor-only mode and to retain the VCF file at the end of the “VariantFiltration” step following hard-filtering as the list of variants for the tumor sample. The second approach was to generate a PoN by pooling together a cohort of normal samples using MuTect and to call mutations with the pooled list of germline variants serving as the normal surrogate^17^. For this, we constructed four panels of normal samples, one for each of our tumor types, using the normal BAM files. The procedure for creating a PoN has been documented previously but in brief, each normal BAM file was passed to the tool separately under the input label of ‘;;tumor’;; with the “artefact_detection_mode” set on^17^. The output vcf per sample was merged together using CombineVariants from the GATK engine to generate the final normal panel.

### Features

We hypothesize that true variants will exhibit different underlying sequencing properties, genomic background characteristics and population distributions when compared to germline variants or sequencing artifacts. Thus to characterize this, we extracted a set of features, based on variant positions found in the final callset, from the VCF file, following GATK processing with HaplotypeCaller, and the tumor BAM file, and annotated these with minor allele and population frequencies from the 1000 Genomes Project and NHLBI^44^ in addition to TCGA recurrences calculated at both the position level and gene level using the latest release of TCGA MAF files. This resulted in a total of 59 features for the PRAD models and 61 features for the BRCA, KIRC and HNSC models. The discrepancy between the total number of features across the four models is a result of different versions of GATK used during processing (ClippingRankSum and SOR are features missing from the PRAD model). The full set of features used is described in **Table 1**.

### Model Training

For our classifier, we chose to use a random forest (RF)^45^ – an ensemble approach that is resistant to outliers and can effectively handle highly-correlated variables. We trained our models using the randomForest package (v4.6-10) in the R statistical environment (v3.2.3). For each tumor type, we set aside 30% of our samples aside as a held-out test set and performed a grid search to parameterize our random forest using our 70% training set in a 10-fold cross-validation scheme (**Figure 1A**). To split samples into 70% training and 30% testing, we rounded 30% of the sample size per tumor type to the nearest integer and set this as the number of samples to allocate for each of our test sets. We then used the base function “sample” from the “stats” package in the R statistical environment (v3.2.3) with the seed 333 to obtain indices to assign alphabetically-sorted sample names to the test set. The samples that remained were assigned to training.

### Random Forest Parameterization

It has been noted that some parameters of random forest are more sensitive to tuning than others^31,46^. In addition, due to the nature of our dataset, a large class imbalance exists between the negative and positive classes (the number of germline SNPs called greatly outweigh the number of somatic SNVs called). Thus to improve our predictive power, we elected to down-sample the major class as a function of the minor class and performed a grid search to tune parameters of random forest using our training set. We assessed the performance using a 10-fold cross-validation scheme to obtain the most optimal values to use for our full model (**Figure 1A**). The three parameters we chose to tune were *mtry, nodesize* and *ntree*.

For the parameter *mtry*, we elected to test factor levels of the default value. Since the nature of our problem is classification and the default *mtry* for classification is the square root of the number of features – which in our case was 7 – we thus used, in addition to the default value of 7, half of the default (4), twice the default (14) and three times the default (21). For *ntree*, we chose to test 1000, 5000 and 10,000 and values of 5, 25, 50 and 100 for the parameter *nodesize*. For all of our datasets, we down-sampled our major class at a ratio of 1:1 with our negative class in order to mitigate bias that can arise from class imbalance.

To split our 70% training set into 10 reasonably equal sets, we took an approach that was similar to the splitting of our training and test sets. We truncated what was 10% of the number of samples in our training set to the nearest integer and set this as the number of samples to allocate per fold. We then used the base function “sample” from the “stats” package in the R statistical environment (v3.2.3) with the seed 99 to to obtain array indices for our vector of sample names. Each time an index was chosen, that was taken out of the pool of indices we may sample from. We repeated this process nine times and used the indices to assign each sample to its respective set, with the remaining samples assigned to fold 10. The last fold also, on average, contained 1-2 more samples than the rest of the folds. For our grid search parameterization, we kept each fold consistent across all parameter combinations tested.

For each parameter combination, we calculated the precision, recall as well as the area under the precision-recall curve (AUPRC). We ranked each parameter combination based on the average area under the precision-recall curve AUPRC across ten folds (ties were broken using the standard deviations where a higher rank was attributed to the lower standard deviation) and chose the set of parameters that generated the highest overall AUPRC. We then trained the full models using *mtry* of 4, *nodesize* of 5 and 10,000 trees.

### Performance Assessment

By varying the vote threshold from 0 to 1, we were able to calculate, across a continuous range, the number of false positive (FP), false negative (FN), true positive (TP) and true negative (TN) calls by using different cutoffs. This was facilitated by the pROC package^47^ (v1.8) in the R statistical environment (v3.2.3). We then used these to calculate metrics for assessing model performance such as sensitivity, specificity and precision. We also constructed a curve using our continuous range of precision and recall values and used the area under this precision-recall curve (AUPRC) as the main metric for comparing model performance. This was done using the trapezoid method and calculated by:

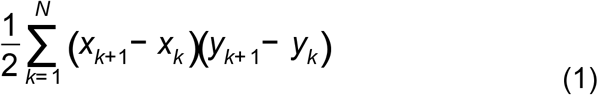

where *x* is the the recall and *y* is the precision at cutoff *k*. The set of parameters with the highest average rank of AUPRC across the ten folds was selected for the full model.

When selecting an operating point for our models based on the AUPRC, we elected to pick a threshold that maximized the harmonic mean of precision and recall, also known as the F1 score, which was calculated by:

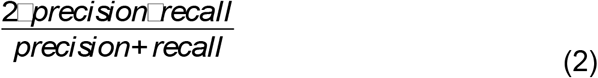

where precision is defined as:

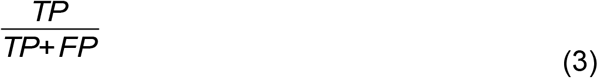

and recall is defined as:

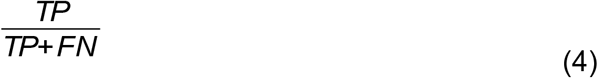

### Cross-tumor and Cross-caller Performance, and Model Convergence

We took three approaches to gauge the generalizability of each model and assessed the effects of both tumor type and SNV calling tool, as well as cohort size on overall performance. First, given models trained using the somatic calls from a single SNV caller (*i.e*. MuTect), we applied the model of one tumor to the test sets of the other tumors and evaluated its cross-tumor performance using the AUPRC metric. Due to discrepancies between features across models, when testing the BRCA, HNSC or KIRC models on the PRAD test set, we manually set the two missing features (ClippingRankSum and SOR) to 0 for all positions in the test set. Alternatively when testing the PRAD model on the BRCA, HNSC or KIRC test set, we omitted those two features from the test data frame when predicting using our RF classifier. Similarly, we assessed cross-caller performance by applying the model trained using one SNV caller and assessed its performance on the other two callers using the AUPRC metric.

Finally, to determine the effects of training size on model performance, we conducted a convergence experiment using reasonably-selected sample sizes in our training set and assessed its performance on the same test set by calculating the AUPRC. We tested sample sizes of 5, 10, 25, 50 and 100 (as well as additional increments of 50 as sample sizes allowed) and compared their AUPRCs to that of the full model.

### Data Visualization

Visualizations were generated in the R statistical environment (v3.2.3) using the lattice (v0.20-33)^48^, latticeExtra (v0.6-28), BPG^49^ (v5.6.8) and VennDiagram (1.6.17) packages^50^. Figures were compiled using LaTeX.

### Benchmarking

To assess how our approach compares to methods developed under similar pretenses, we applied three published tools (EBFilter, VNC and VarDict)^18,19,30^ to each of the held-out test sets. We selected these tools based on their proposed functionality and the fact that they accepted similar inputs and generated comparable outputs to our method. We used the same AUPRC metric to assess performance across the different algorithms. Since each tool had its own specific method or score for annotating potential true variants or variants of interest, to generate a curve for each, we varied the threshold for each tool across its full range of values for the entire cohort. All tools were run with default options or with those suggested in the user guidelines for the tool unless otherwise specified.

VarDict was executable given a VCF file, BAM file and the human genome reference fasta as inputs, however, some of the other tools required additional input files. Both EBFilter and VNC took an approach that was similar to MuTect in that both required an additional list of normal samples to create a surrogate normal. For EBFilter, we used the normal samples from the training set of each tumor type to generate this panel in all of the EBFilter runs. For VNC, we downloaded all available (n = 427) variant files of normal samples that were processed by Complete Genomics (CG) for the 1000Genomes Project from the FTP site (ftp://ftp.1000genomes.ebi.ac.uk/vol1/ftp/phase3/data/). We used these samples instead of TCGA normal samples because the tool was designed to work with CG outputs and only accepted the varfile format as input to generate the virtual normal. To generate a range of precision and recall values for EBFilter, we varied the score that was outputted by the tool from 0 to the maximum score of the cohort. Since VNC implemented two fields for filtering calls – both relating to the number of samples out of the total pool of samples used in the construction of the virtual normal in which a variant was found – we varied both thresholds from 0 to 427 and calculated precision and recall at each combination of thresholds. In single-sample mode, VarDict implements a single allele frequency (AF) filter during variant identification, so to calculate precision and recall for the PR-curve, we varied the AF from 0 to 1. When selecting operating points for each tool, we used the same approach as with S22S and chose that which maximized the F1-score. All raw and processed files generated in benchmarking the three tools can be found in **Supplementary Data 1**.

### Availability

Software and models are available at http://labs.oicr.on.ca/Boutros-lab/software/s22s.

